# Decision Making by *Drosophila* Flies

**DOI:** 10.1101/045666

**Authors:** Julius Adler, Lar L. Vang

## Abstract

> “*Decision making has all the secrets of everything: who we are, what we do, how we navigate the world*.” “How Do I Decide? The Brain with David Eagleman”, 2015.

When presented with attractant (light) together with an amount of repellent (methyl eugenol) that exceeds attractant, *Drosophila melanogaster* fruit flies are of course repelled, but nine mutants have now been isolated that were not repelled. Although able to respond to attractant alone and to repellent alone, these mutants fail to make a decision when the two are together during the first two months of the study. They are considered defective in a decision-making mechanism. The defect occurs at 34°C but not at room temperature, so these are conditional mutants. Efforts at genetic mapping have been made. Our aim is to discover how decision making gets accomplished and how this results in a behavioral response. We indicate that there is a mechanistic relationship between decision making and the central complex in *Drosophila* and between decision making and the prefrontal cortex in humans and other vertebrates.

Over a period of six months these mutants changed into ones that are attracted when presented with attractant together with what was overpowering repellent before. Nearly full attraction was achieved at fifteen to thirty days. With attractant alone these mutants were attracted like the original parent and with repellents alone they were repelled like the original parent. The mutants have been genetically mapped.

## I. INTRODUCTION

Decision making occurs in every organism. Decision making – see Fig. 1 – can be between different attractants or between different repellents or between an attractant plus a repellent. Here we consider the last of these. It is the mechanism of decision making that is our interest here.

**Fig 1.**
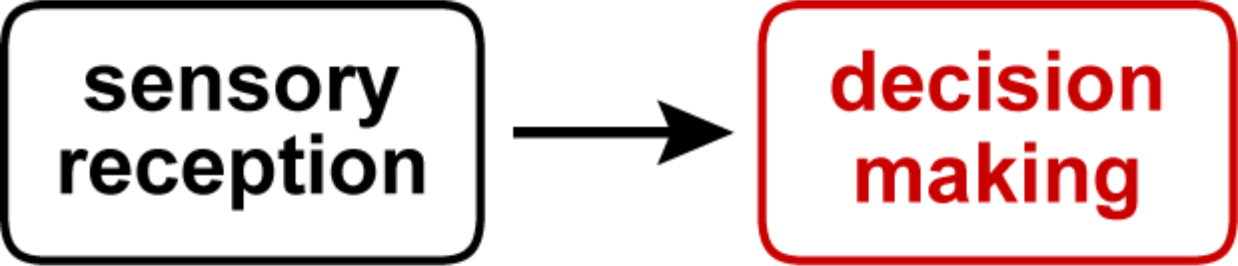
Relation between sensory receptors and decision making. Receptors act on decision making (in red), which is studied here. Information then goes on to the central complex in insects or the prefrontal cortex plus basal ganglia in vertebrates. This result is then sent on by means of motor neurons and muscles to generate a behavioral response.

What an organism does when presented with attractant plus repellent has been studied previously: in the prefrontal cortex of humans and other primates [1–12] and of rodents [13–17]; in other vertebrates as in *The Merging of the Senses* [18], *The Handbook of Multisensory Processes* [19], and *The New Handbook of Multisensory Processing* [20]; in invertebrates [9, 18, 21–26] and *Decision Making in Invertebrates* [27]; in green plants [27]; and in bacteria [29, 30].

In this report we study repulsion of *Drosophila* flies when an attractant, light, is exceeded by a repellent, methyl eugenol. (Other repellents, like benzaldehyde, were tried also, see Supplemental Methods.) We report here the isolation and properties of mutants that fail in this repulsion though they are motile and respond normally to attractant alone and to repellent alone, so-called “decision mutants”. The mutants presumably are defective in decision making. In another paper we report *Drosophila* mutants that have a decreased responsiveness to many different stimuli [31].

## II. RESULTS

### A. ASSAYING THE BEHAVIOR

The parent and mutants were assayed as shown in Fig 2. At one end was attractant (light) together with overpowering repellent (methyl eugenol). At the other end the flies started out. The parent stayed largely where placed on account of the overpowering repellent but motile mutants were found that failed to stay there.

**Fig 2.**
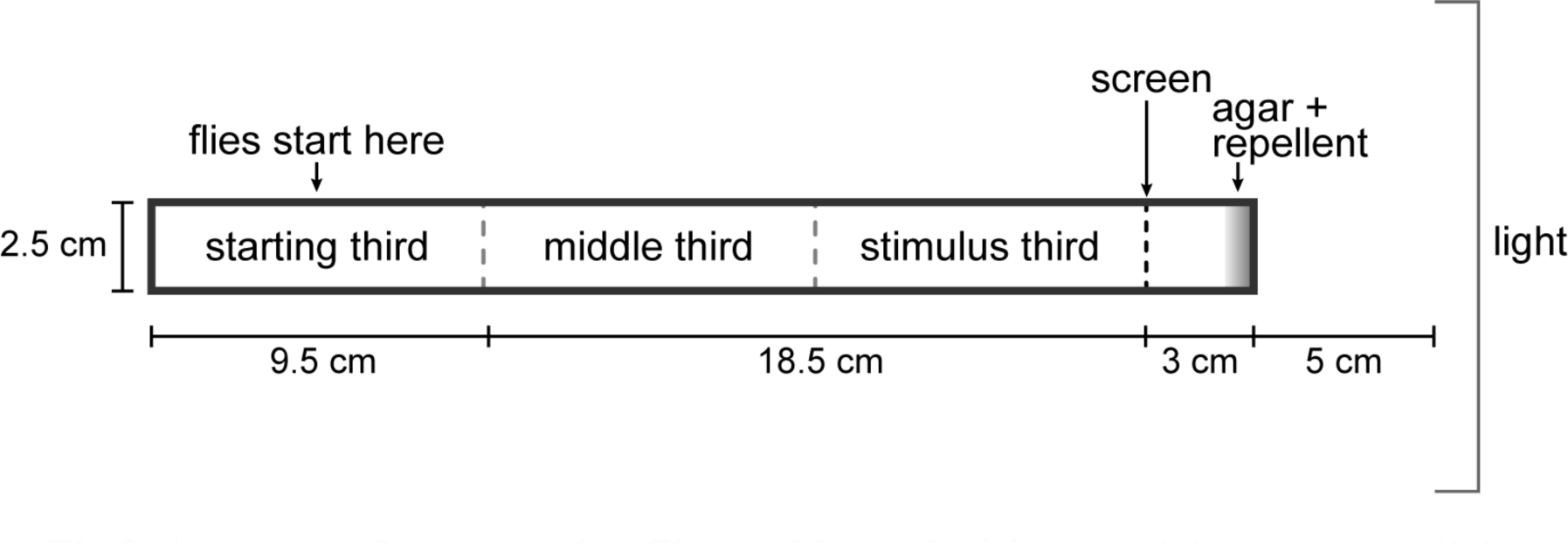
Apparatus for measuring flies making a decision. At right, attractant (light, 4000 lux) together with overpowering repellent (0.1M methyl eugenol), at left flies start out. Parent stays where put, mutants do not. When attractant only is used, the setup remains the same except there is no repellent. When repellent only is used, the setup remains the same except the light source is placed parallel to the setup. See details in IV.

Several kinds of motile mutants might be expected: 1) Mutants that are able to sense the attractant better than the parent and therefore they are attracted in spite of presence of this amount of repellent. These would probably be mutants in the attractant receptor pathway. 2) Mutants that are able to sense the attractant normally but sense repellent less well than the parent. These are attracted and would probably be mutants in the repellent receptor pathway. 3) Mutants with unchanged sensing of attractant alone and unchanged sensing of repellent alone, but they are not repelled when attractant is together with overpowering repellent, perhaps due to a defect in decision making. It is this third kind that we were eager to find.

Nine flies were isolated: mutants 2, 3, 5, 6, 7, 8, E, I, and J. In the first 10 days after eclosion, during the first two months, they and their parent were tested in the apparatus of Fig 2 at 34 and 21°C for response to attractant together with overpowering repellent, then response to attractant alone and response to repellent alone. See a preliminary report [32].

### B. RESPONSE OF PARENT AND MUTANTS TO ATTRACTANT TOGETHER WITH REPELLENT AT 34°C, EARLIER AGE

The response of parent and mutants to attractant (light) together with overpowering repellent (methyl eugenol) at 34°C was studied in the first 10 days after eclosion, during the initial two months. As shown for the parent in Fig 3A, the repellent overcame the attractant when his amount of attractant and this amount of repellent were used. In the mutants that was not the case, for example see Fig 3B for mutant 2; the mutant flies were more or less randomly distributed throughout the tube at 34°C. Fig 3C shows results for parent and all nine of the mutants at 34°C in the first 10 days after eclosion, during the first two months. While the parent was repelled, the mutants were largely not repelled.

**Fig 3.**
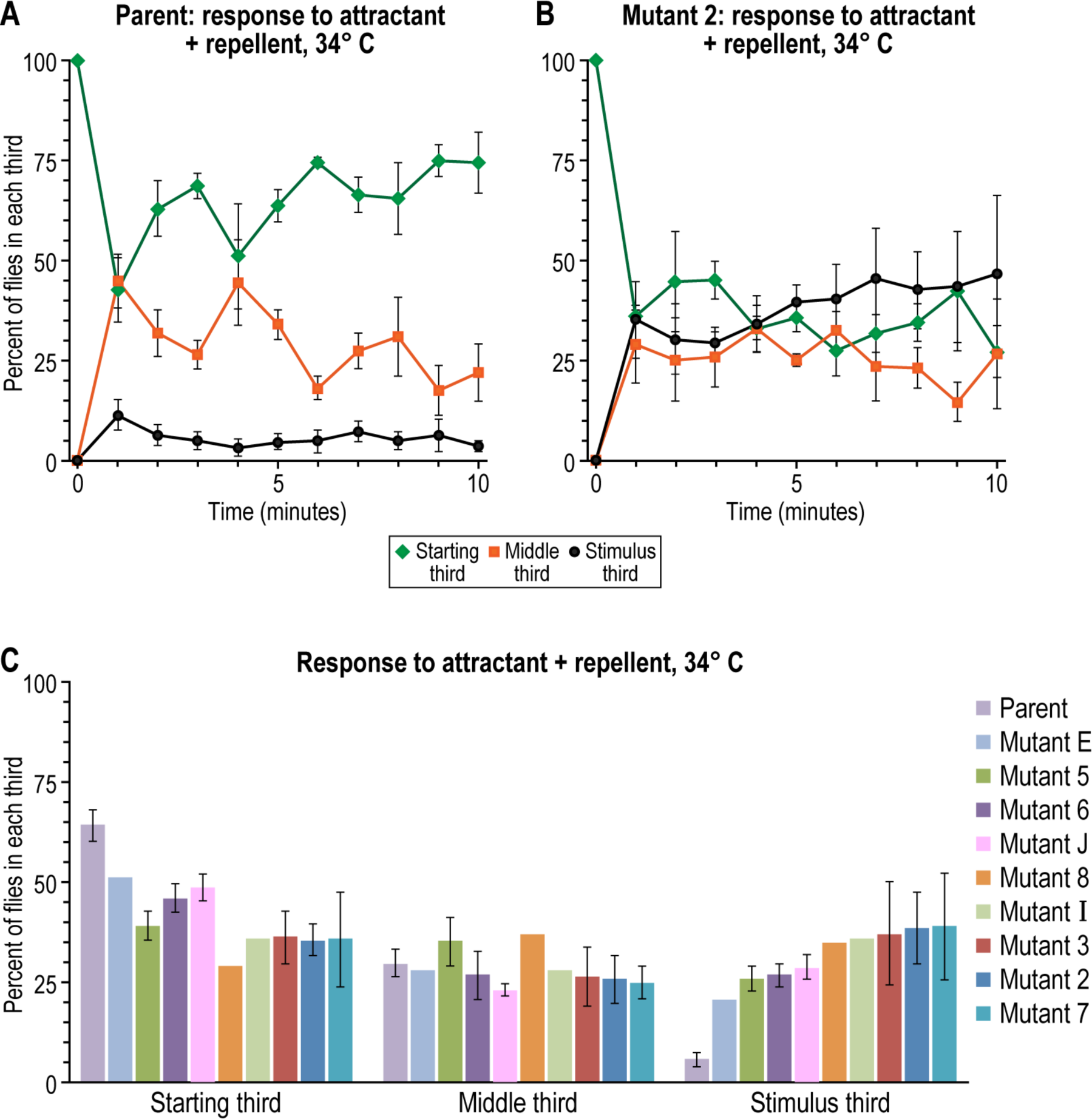
Response to attractant (light, 4000 lux) together with repellent (0.1M methyl eugenol) by parent and mutants at 34° C. (A) Parent, time course over 10 minutes; the repellent overcame the attractant. (B) Mutant 2, time course over 10 minutes; the repellent did not overcome the attractant. (C) Parent and all the mutants, average of each 10 minute time course (n=4, 1, 3, 3, 12, 1, 1, 3, 3, 4 for the parent, E, 5, 6, J, 8, I, 3, 2, and 7, respectively; mean±SEM). Data without error bars: experiments were done only once; see section G. With mutant J, 1000 lux light and 0.3M methyl eugenol were used. Approximately 10 to 20 flies were used per trial.

### C. RESPONSE OF PARENT AND MUTANTS TO ATTRACTANT TOGETHER WITH REPELLENT AT ROOM TEMEPRATURE, EARLIER AGE

All those experiments were carried out at 34°C, but at room temperature (21°C) the parent and early mutants looked alike in the first 10 days after eclosion, during the
first two months, see Fig 4.

**Fig 4.**
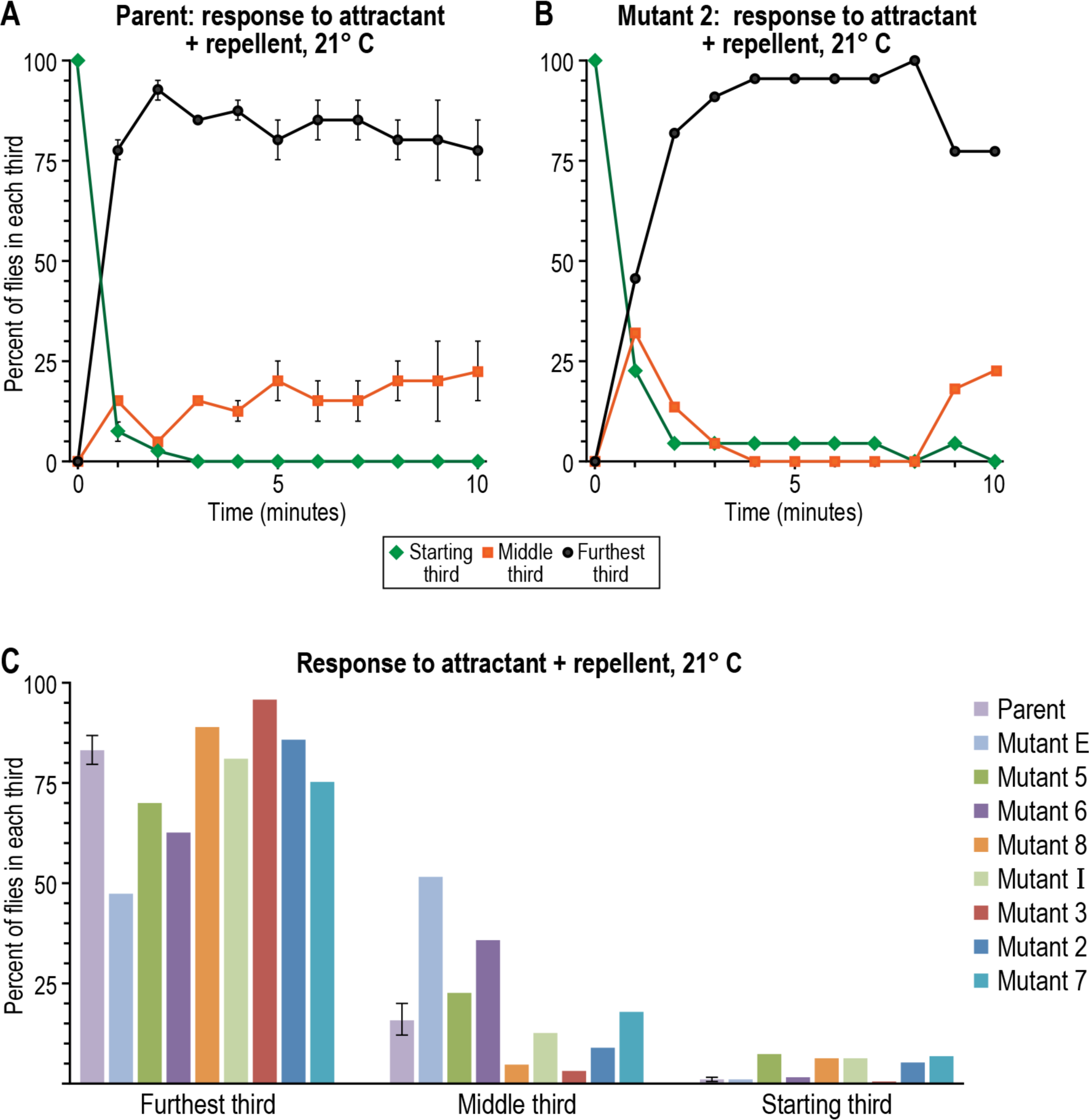
Response to attractant (light, 4000 lux) together with repellent (0.1M methyl eugenol) by parent and mutants at 21°C. (A) Parent, time course over 10 minutes (n=2, mean±SEM). (B) Mutant 2, time course over 10 minutes (n=1). (C) Parent and all the mutants, average of each 10 minute time course (n=2 for parent, n=1 for mutants E, 5, 6, 8, I, 3, 2, and 7). Data without error bars: experiments were done only once; see section G. Note that in the experiments of Fig 4 the flies started out at the end closest to attractant plus repellent (unlike in Fig 3, 5, and 6, where they started out at the end away from attractant plus repellent) before they went to the end away from attractant plus repellent. Approximately 10 to 20 flies were used per trial.

Therefore we conclude that these mutants are conditional, i.e. the defects show up at the higher temperature, but not at room temperature.

### D. RESPONSE OF PARENT AND MUTANTS TO ONLY ATTRACTANT AT 34°C, EARLIER AGE

The parent and each of the early mutants were assayed at 34°C with only attractant (light) presented in the first 10 days after eclosion, during the first two months. The result was that the responses were essentially the same for parent and mutants, see Fig 5.

**Fig 5.**
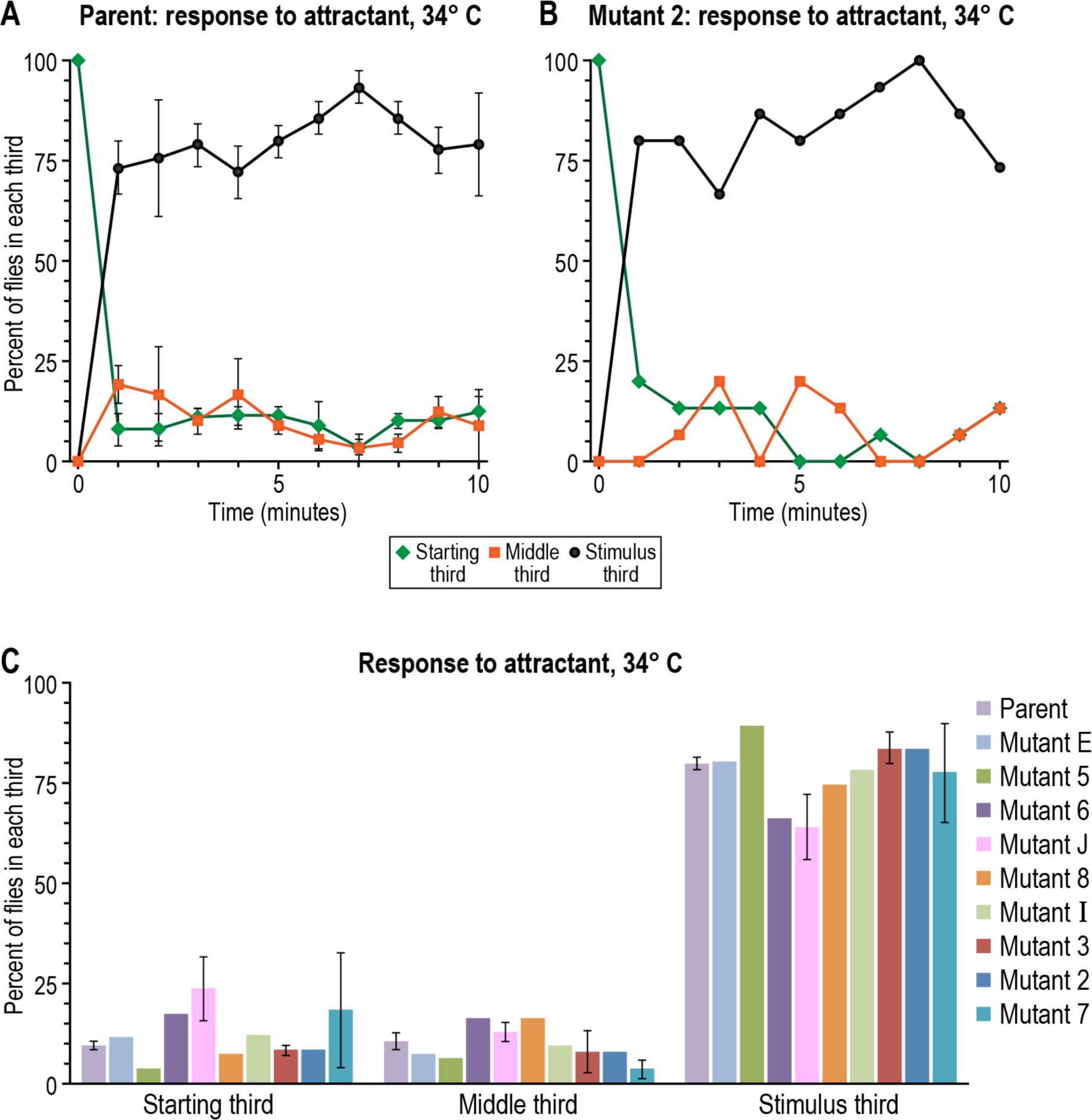
Response to attractant (light, 4000 lux except 1000 lux for Mutant J) by parent and mutants at 34° C. (A) Parent, time course over 10 minutes (n=4, mean±SEM). (B) Mutant 2, time course over 10 minutes (n=1). (C) Parent and all the mutants, average of each 10 minute time course (n=4, 1, 1, 1, 10, 1, 1, 2, 1, and 2 for parent, E, 5, 6, J, 8, I, 3, 2, and 7, respectively; mean±SEM). Data without error bars: experiments were done only once; see section G. Approximately 10 to 20 flies were used per trial.

### E. RESPONSE OF PARENT AND MUTANTS TO ONLY REPELLENT AT 34°C, EARLIER AGE

With repellent (methyl eugenol) only, responses were essentially the same for parent and mutants in the first 10 days after eclosion during the first two months, see Fig 6.

**Fig 6.**
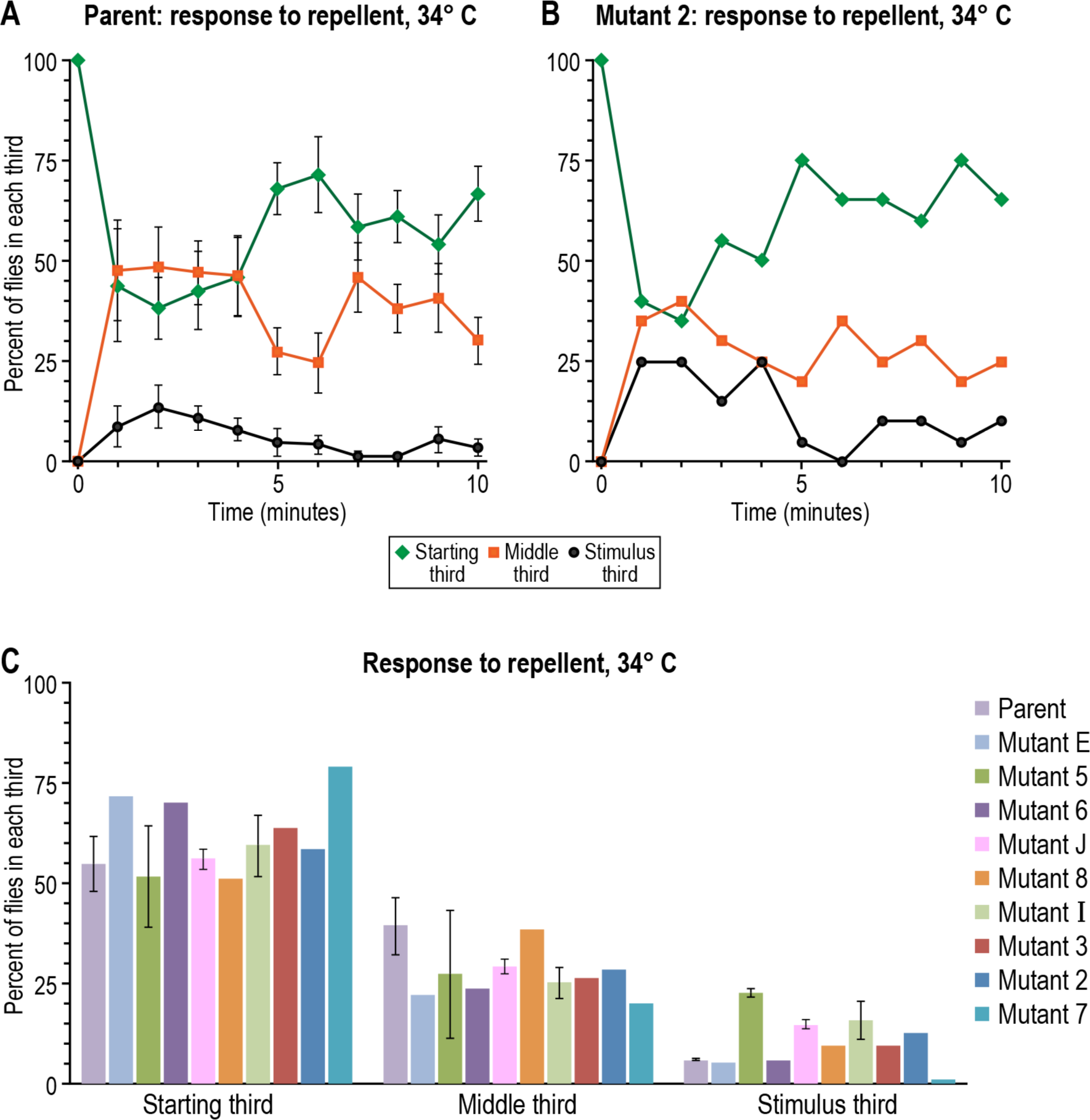
Response to repellent (0.1M methyl eugenol) by parent and mutants at 34° C. (A) Parent, time course over 10 minutes (n=4, mean±SEM). (B) Mutant 2, time course over 10 minutes (n=1). (C) Parent and all the mutants, average of 10 minutes (n=4, 1, 2, 1, 10, 1, 3, 1, 1, and 1 for parent, E, 5, 6, J, 8, I, 3, 2, and 7, respectively; mean±SEM). Data without error bars: experiments were done only once; see section G. With Mutant J, 0.3M methyl eugenol was used. Approximately 10 to 20 flies were used per trial.

Since the mutants failed to give a mutant response when only attractant or only repellent was used, their defect appears to be not in the attractant receptors and not in the repellent receptors but in a place where the action of attractant and the action of repellent come together, i.e. in decision making.

### F. STABILITY OF THE EARLY MUTANTS AND MAPPING THEM

The early mutants described here have limited stability. One of them, mutant J, died at about six months after isolation. The rest of them stayed alive but by six months after first becoming adults they had changed their property: they are now attracted by attractant plus what was previously overpowering repellent (see below), and they now show this attraction up two or three weeks after eclosion instead of immediately after eclosion. We report on the property of these later mutants below.

Due to the above changes in the mutants, we were not able in time to carry out more than one test for some of the data in Fig 3C, 4B and 4C, 5B and 5C, and 6B and 6C. Also for this reason, we did not succeed in time to finish genetically mapping the mutants. However, based on preliminary recombination mapping data using the *y cho cv v f* chromosome, we showed that mutant 2 falls within 11F to 15F of the X chromosome (data not shown).

### G. RESPONSE OF PARENT AND MUTANTS TO ATTRACTANT TOGETHERWITH REPELLENT AT 34°C, LATER AGE

Six later mutant flies were studied here: 2a, 6a, 7a, 8a, Ea, and Ia. “Later mutants” are descended from mutants 2, 6, 7, 8, E, and I, respectively, which had been studied above at an earlier age. These six and their parent were now tested in the first 10 days after eclosion, one year after their initial isolation, in the same way as above for response to attractant together with overpowering repellent, then response to attractant alone and response to repellent alone.

The parent (Fig 7A) was strongly repelled by attractant plus overpowering repellent in the first week after eclosion. Then gradually it was repelled less but still it was largely repelled by 20 days (Fig 7A). In short, repulsion was strong but it became less with age.

**Fig 7.**
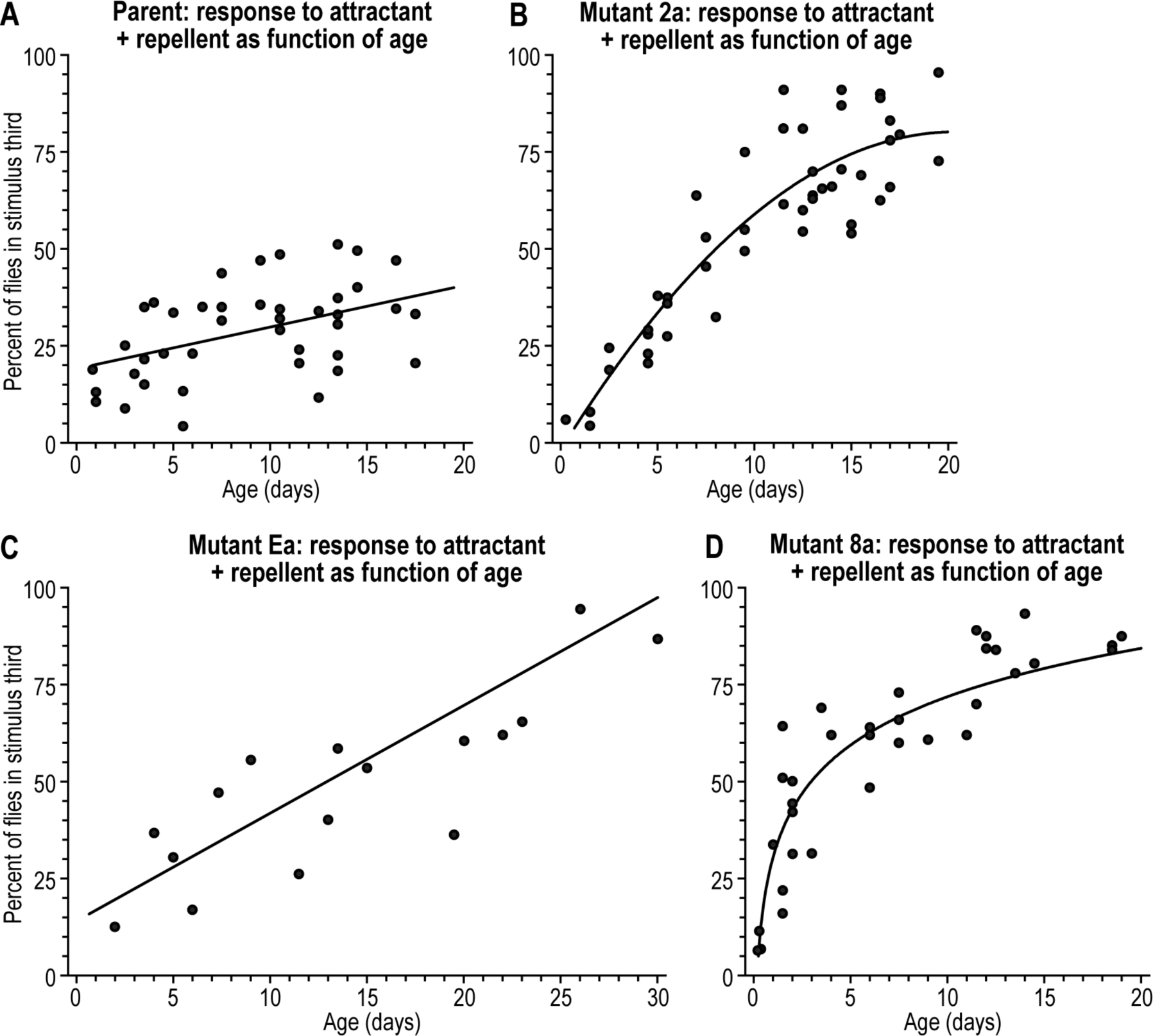
Response to attractant (light, 4000 lux) together with repellent (methyl eugenol, 0.1M) at 34°C as function of age after eclosion. Whereas the parent (A) shows weak attraction over time, the mutants (B to D) are attracted at a faster rate. In each case this figure shows the stimulus third as function of age; see S7 Fig for all three thirds. Approximately 10 to 20 flies were used per trial.

Mutant 2a (Fig 7B) started out repelled by attractant plus overpowering repellent, like the parent, then it was repelled less, giving about half attraction at 10 days, and then it gave close to full attraction at 15 to 20 days. For a comparison of parent and Mutant 2a with data combined from 12 to 19 days of age, see Fig 8A and 8B. The response of Mutant 7a was much like the response of Mutant 2a in Fig 7B, taking about 20 days for nearly full attraction to occur (data not shown).

**Fig 8.**
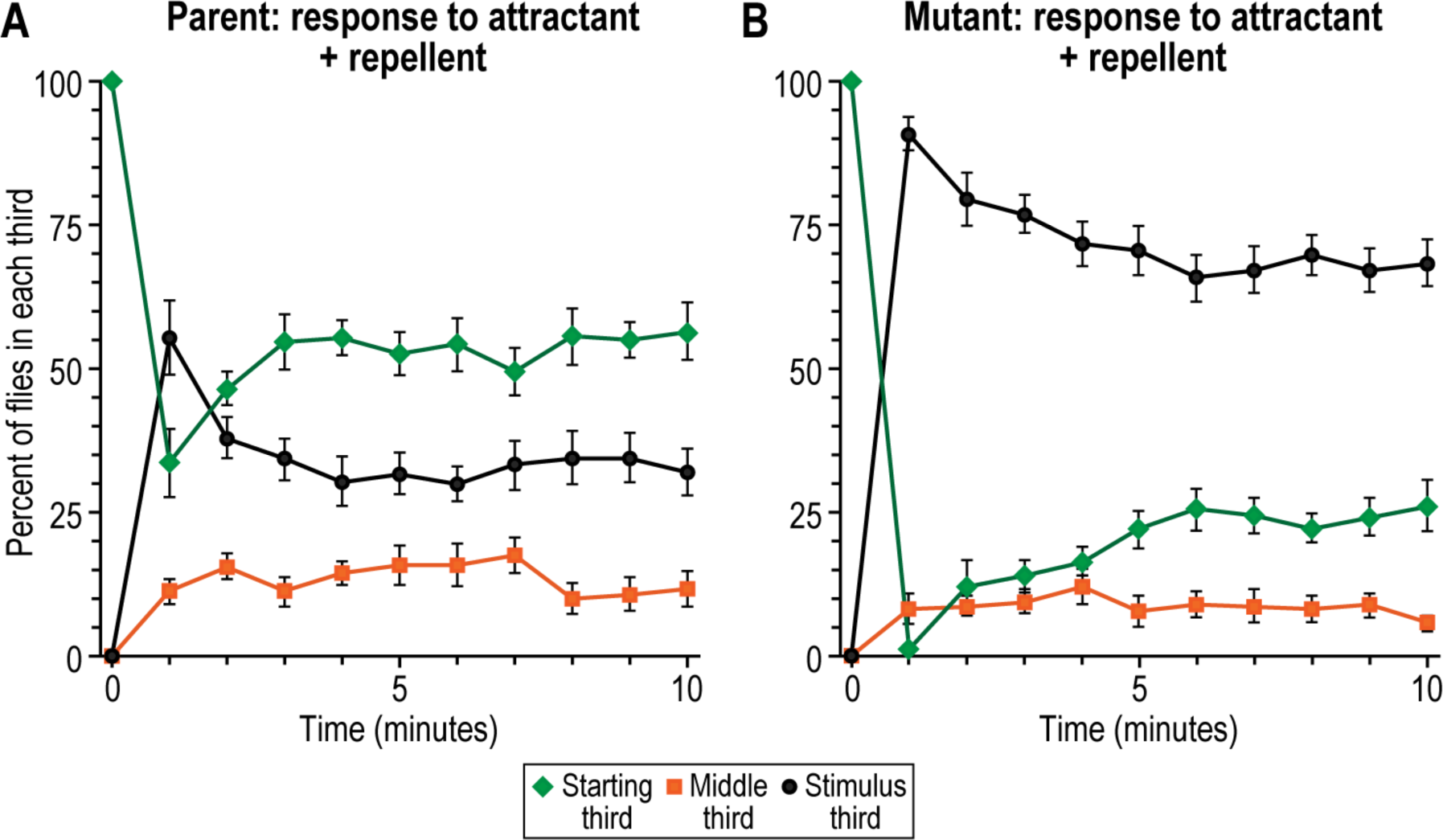
Response to attractant (light, 4000 lux) together with repellent (methyl eugenol, 0.1M) at 34°C by parent and mutant. (A) Most of the parent stayed largely in the starting third because of the overpowering repellent (green curve). (B) Mutant 2a went to the attractant (black curve) in spite of this repellent. Data are combined from 12-19 day old flies. Data are mean±SEM. Approximately 10 to 20 flies were used per trial.

Mutant Ea (Fig 7C) also started out repelled, reached half of full attraction at about 13 days, then full attraction at about 30 days. (This mutant was not studied further.) The response of Mutant Ia was much like that of Mutant Ea, taking about 30 days for full attraction to occur (data not shown).

Mutant 8a (Fig 7D) also started out repelled, reached half of full attraction by about 3 days, then close to full attraction by day 14-15. Mutant 6a gave responses similar to Mutant 8a (data not shown).

In summary, the attraction of these six mutants is slow to develop when both attractant and more repellent are present together, but eventually it amounts to nearly complete attraction.

How much attractant it took to overcome repellent is reported in Fig 9. For the parent (Fig 9A) repulsion by 0.1M methyl eugenol failed to be overcome or was overcome only weakly by any of the light intensities studied from 0 to 10,000 lux. For Mutant 2a (Fig 9B) and for Mutant 8a (Fig 9C) repulsion by 0.1M methyl eugenol was overcome by light at about 1,000 lux. Thus for Mutants 2a and 8a, unlike for the parent, the attractant almost completely took over when its intensity was raised sufficiently over that of repellent.

**Fig 9.**
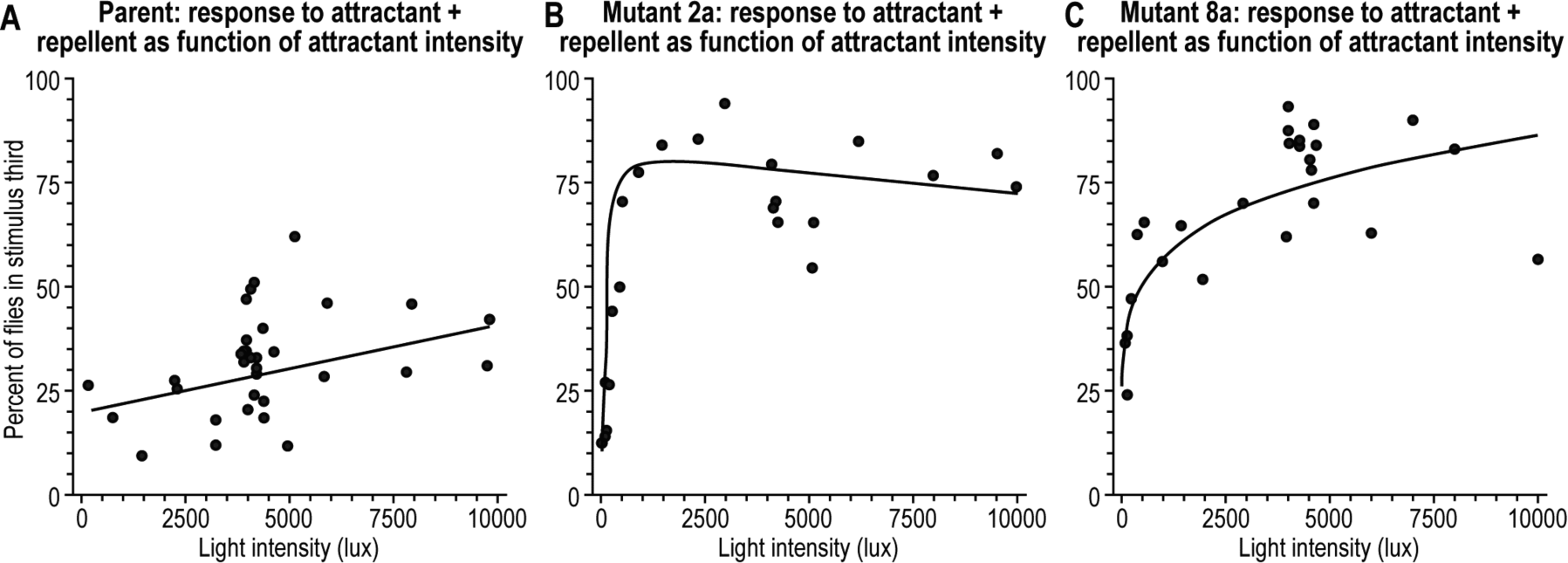
Response to attractant (light,) together with repellent (methyl eugenol, 0.1M) at 34°C as function of attractant intensity. In each case this figure shows the stimulus third; see S9 Fig for all three thirds. Approximately 10 to 20 flies were used per trial. (Parent age: 12 to 20 days; Mutant 2a age: 10 to 19 days; Mutant 8a: 11 to 18 days.)

### H. RESPONSE OF PARENT AND MUTANT TO ATTRACTANT TOGETHER WITH REPELLENT AT ROOM TEMPERATURE, LATER AGE

All the above experiments were carried out at 34°C. Next are experiments done at room temperature (21° to 23°C). Since diffusion of repellent is slower at room temperature than at 34°C, 30 minutes instead of the usual 15 were allowed for the repellent to diffuse sufficiently before the measurements began. The flies were started at the end that had attractant (light) together with repellent (0.06M methyl eugenol), from which they ran away. The results were the same for parent and mutant 2a (the only mutant tested at room temperature) at the age of 8 to 19 days. For parent the result was 13, 39, 48% (n=17) and for Mutant 2a it was 18, 36, and 46% (n=6), where the first of the three numbers represents the third closest to attractant plus repellent; the second, the middle third; and the third, the part furthest away from attractant plus repellent. Thus the difference between parent and mutant found at 34°C did not occur at room temperature. That makes the mutant a conditional one. Once the action of that gene at 34°C is discovered, one can try to understand the need for conditionality here. Recall that a conditional response was also found for the early mutants.

### I. RESPONSE OF PARENT AND MUTANTS TO ONLY ATTRACTANT AT 34°C, LATER AGE

The response of parent and later mutants to only attractant (light) at 34°C as a function of age after eclosion is shown in Fig 10. At first there was a period of no attraction, then there was attraction to the light by parent (Fig 10A) and by Mutants 2a (Fig 10B) and 8a (Fig 10C). The curves are not identical, though they are close.

**Fig 10.**
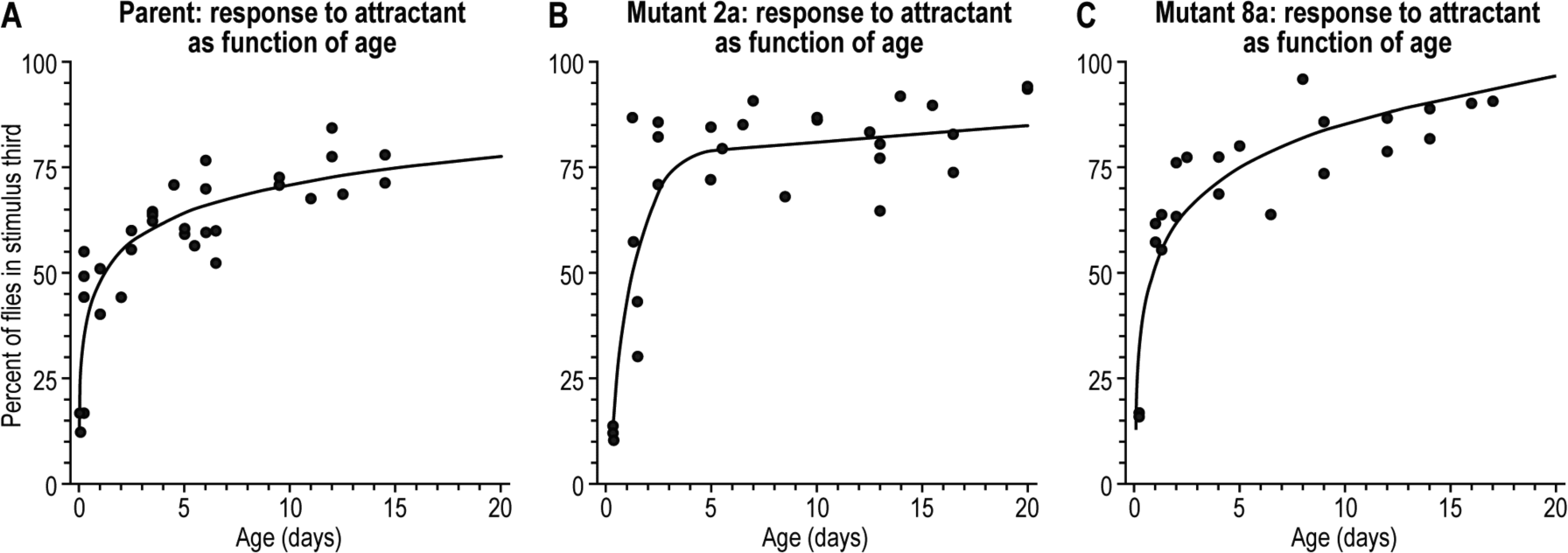
Attraction to light (light, 4000 lux) at 34°C as function of age after eclosion. In each case this figure shows the stimulus third as function of age; see S10 Fig for all three thirds. Approximately 10 to 20 flies were used per trial.

Notice that with only attractant (Fig 10B) the attraction occurs faster than with attractant + repellent (Fig 7B). This shows that the repellent does have an effect in Fig 7B: it slows the attraction down. One might otherwise have argued that in Fig 7B the result is due only to attraction, i.e. that the repellent is not at all operating here.

The response as a function of attractant intensity at 34°C was compared for parent (Fig 11A) and Mutants 2a (Fig 11B) and 8a (Fig 11C). Up to about 4,000 lux the curves were similar, but at higher attractant intensity (5,000 to 10,000 lux) Mutant 8a was different: 10,000 lux is as high as 1,000 lux for Mutant 8a but not for parent and Mutant 2a.

**Fig 11.**
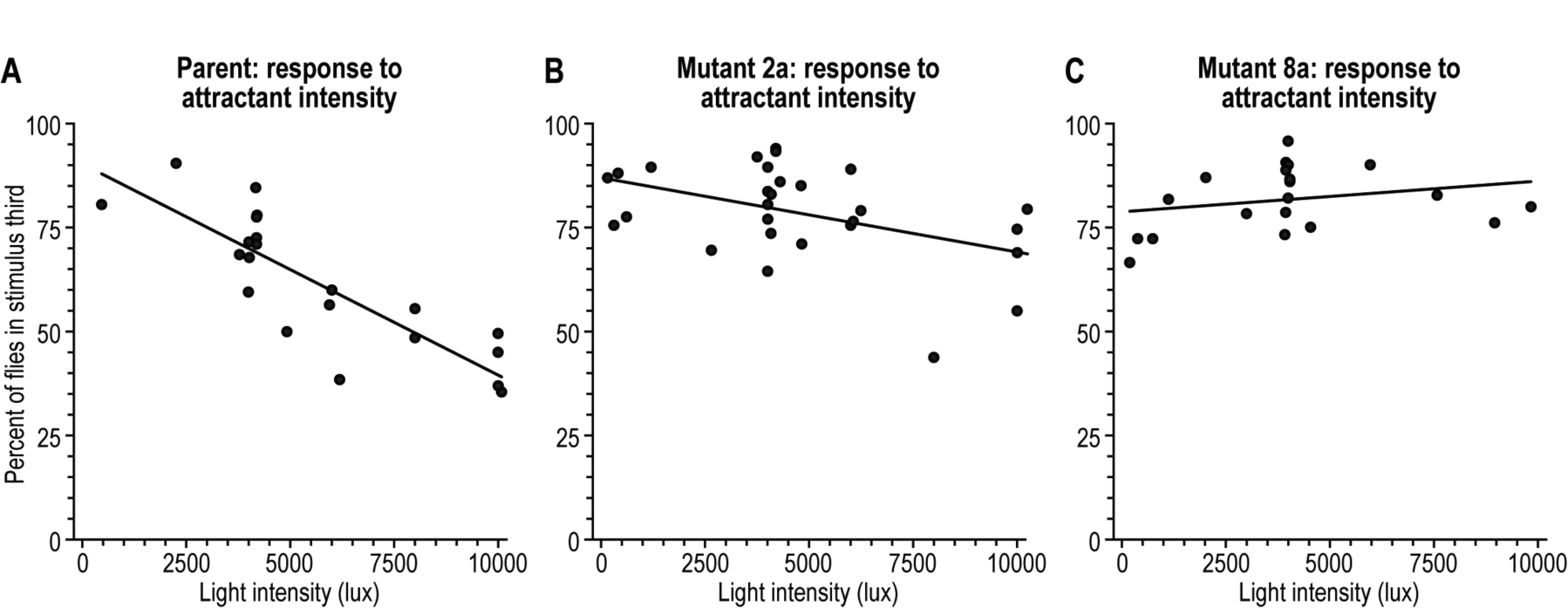
Response to attractant at 34°C as function of attractant intensity. In each case this figure shows the stimulus third; see S11 Fig for all three thirds. Approximately 10 to 20 flies were used per trial. (Parent age: 10 to 17 days; Mutant 2a age: 10 to 20 days; Mutant 8a age: 8 to 17 days.)

### J. RESPONSE OF PARENT AND MUTANTS TO ONLY REPELLENT AT 34°C, LATER AGE

The response of parent and mutants 2a and 8a to repellent (0.1M methyl eugenol) as a function of age after eclosion is shown in Fig 12 over a 20 day period for the third closest to repellent. The three strains were repelled about equally over a 20 day period. The responses as a function of repellent concentration (10^−3^, 10^−2^, and 10^−1^ M methyl eugenol) at 34°C were similar for parent and Mutants 2a and 8a (shown in S13 Fig), so again the three strains were alike in that regard.

**Fig 12.**
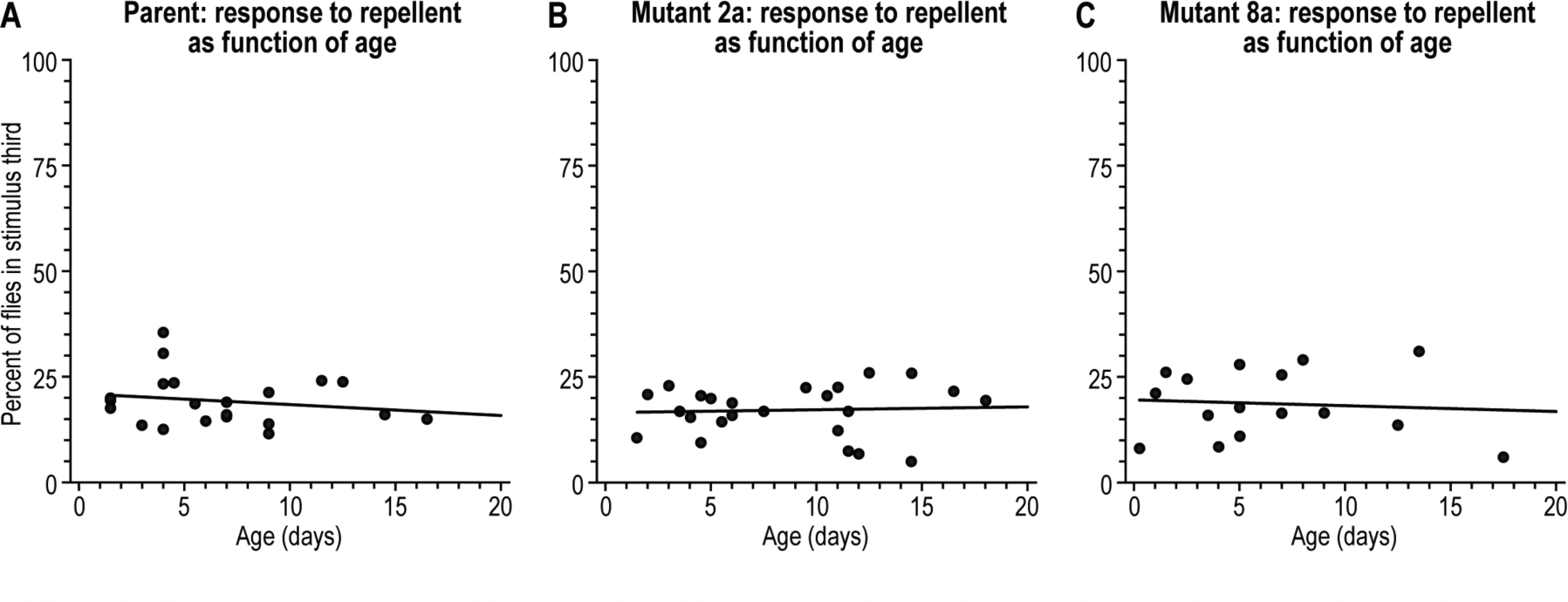
Response to repellent at 34°C as function of age after eclosion. In each case (parent, mutants 2a and 8a) the stimulus third was repelled about equally by 0.1M methyl eugenol over a 20 day period. See S12 Fig for all three thirds. Approximately 10 to 20 flies were used per trial.

### K. MAPPING THE LATER MUTANTS AT 34°C

By use of genetic mapping we have determined where some of the present mutants are located on the fly’s chromosomes. Using deficiency mapping, we found that *Df(1)BSC766* uncovered Mutant 2a. Of the genes in this region, *Syt12* failed to compliment. This gene has been previously shown to be involved in neurotransmitter secretion [33]. We have not yet been able to localize Mutant 8a to a specific locus. So far we predict that the mutants described here may map in the same genes as the early mutants described above because the effects appear to be related (see Fig 13).

## III. DISCUSSION

### A. EARLY FLIES

When parental *Drosophila* flies were presented with an attractant (light) together with a higher amount of repellent (methyl eugenol) they were of course repelled (Fig 3A). Here we report the isolation and study of *Drosophila* mutants that are neither repelled nor attracted when they are presented with attractant together with a higher amount of repellent (Fig 3B and 3C), although they are attracted by attractant alone (Fig 5) and repelled by repellent alone (Fig 6), so apparently they fail just in making a decision. This is summarized in the green of Fig 13.

**Fig 13.**
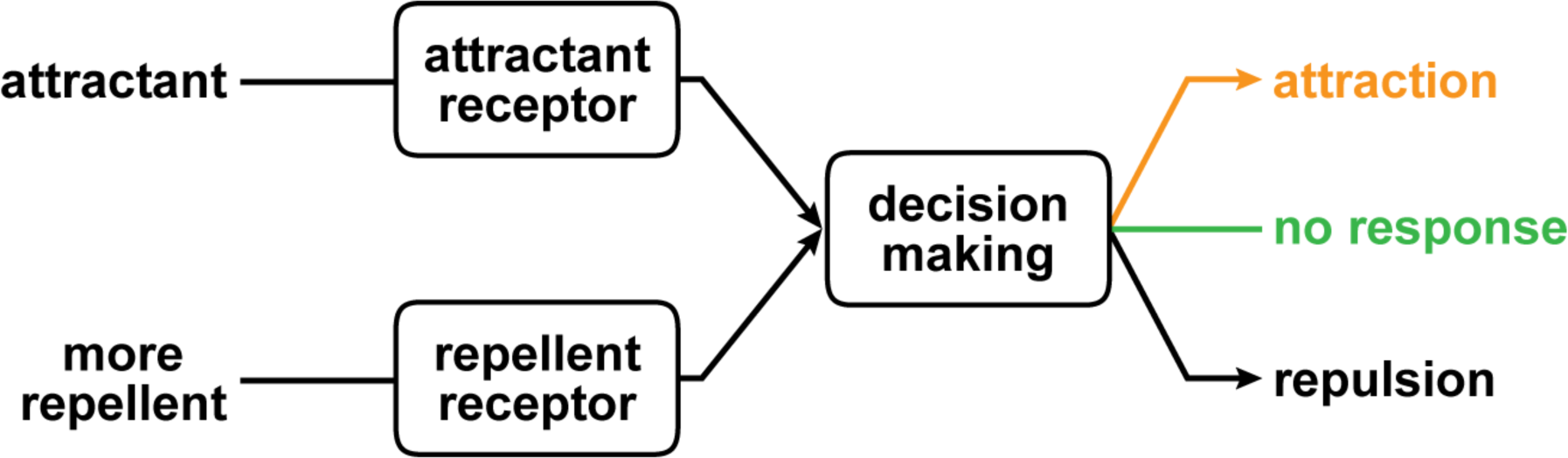
A model of decision making. Parental flies presented with more repellent than attractant are repelled (see repulsion in black at right). Early mutants fail to respond (in green), presumably due to a defect in decision making. Later mutants are attracted (in
orange) due presumably to a different defect in decision making (see below).

What is defective in these mutants? The interaction between stimuli and decision making has been studied/reported by Marcus Raichel [34], Dennis Bray [35], and Björn Brembs [36–38]. There is an active state that is present before stimuli are presented, then a different state upon presentation of stimuli. Although discovered in mammals [34], such a phenomenon occurs also in invertebrates and bacteria [35].

Where does the consequence of these mutations take place? Some possibilities are in the mushroom bodies [39–46] or in the central complex [47–49]. The central complex consists of the protocerebral bridge, the fan-shaped body, the ellipsoid body, and noduli [50, 51].

Several approaches can be used to better elucidate the entire mechanism starting from decision making to the behavioral response. One is to start from decision-making mutants like those obtained here and go step-by-step in finding mutants for the remaining genes all the way to the behavioral response. The second approach is to test central complex mutants already isolated by others [50–57]. These were kindly given to us by Roland Strauss, Burkhard Poeck, and Douglas Armstrong. They were tested here for ability to block the response of flies to attractant plus overpowering repellent. We found that *cex*^*ks181*^, *cbd*^*ks96*^, *ebo*^*ks267*^ [55] and *GAL4-210y/UAS-shi*^*ts1*^ [51] had a similar response to the wild-type when they were tested with attractant (light, 1000 lux) alone and overpowering repellent (methyl eugenol, 0.1M) alone but they had a mutant response when attractant (light, 1000 lux) was presented together with the overpowering repellent (methyl eugenol, 0.1M) (S15 Fig).

In work by others [39–46], *Drosophila* that were flying (but ours were walking) were exposed to attractive or repulsive odor in addition to the visual stimuli encountered in flight. Integration of olfactory and visual information appeared to take place in neurons leading to muscles [44, 46].

### B. LATER FLIES

We report mutants, actually derived from those early ones, that decided in favor of attractant even though the same-as-before higher amount of repellent was presented (Fig 7B, 7C, and 7D) but yet they responded normally to attractant alone (Fig 10) and to repellent alone (Fig 12). Fig 13, in orange, tells a summary of these responses to attractant plus the same-as-before higher amount of repellent.

The curve for Mutant 2a (Fig 7B) goes up much faster than the curve for the parent, which does go up slowly (Fig 7A). Thus attraction overcomes repulsion faster in the mutant than in the parent. What is that process affecting the balance between repulsion and attraction and how has it been changed by mutation in 2a? This has to be determined in order to understand what is wrong in the mutant. Finding that out should identify a factor in decision making.

Mutant 2a and its parent look much alike when presented with attractant alone (Fig 10) or repellent alone (Fig 12). So Mutant 2a is normal when encountering single stimuli but abnormal when there is a choice to be made (Fig 7B). Thus Mutant 2A appears to be a decision mutant. Perhaps the same is true for Mutants 7a, Ea, and Ia, but they have not been tested further.

Mutant 2a gives a response at an early age when presented with only attractant (Fig 10B) but it gives a response at a later age when presented with attractant plus repellent (Fig 7B and 8B). Why is the attractant-plus-repellent response of mutant 2a late to occur? Investigating this may contribute to understanding the mechanism of decision making.

Mutant 8a has higher light sensing ability (Fig 11C) than Mutant 2a (Fig 11B). When attractant is together with repellent, that may explain why light wins out easier for Mutant 8a (Fig 7D) than for Mutant 2a (Fig 7B).

Mutants 2a (Figure 7B), 7a, Ea (Fig 7C), Ia, 8a (Fig 7D) and 6a all show a defect later in the life of the fly (13 to 30 days) than did their precursors (1 to 10 days) (Fig 3). Perhaps the mutants in this report could be considered similar to people who have a problem that occurs at a later age, for example a gradual loss of memory [58, 59] or Alzheimer disease [60].

*Drosophila* has emerged as an ideal model to study such age-dependent neurodegenerative diseases. Early work identified a series of mutant genes involved in degeneration of the brain [61, 62]. Since then, other *Drosophila* genes have been identified to be involved in maintenance of the brain [63, 64].

### C. RELATION OF THE CENTRAL COMPLEX TO THE PREFRONTAL CORTEX

Starting in the 1870’s, it became apparent to some psychologists that there may be a part of the human brain, the prefrontal cortex, that is master of the whole brain [65, 66]. The prefrontal cortex is also known as the “central executive” [67], the “executive brain” [68], and “executive control” [69]. The prefrontal cortex in people and other primates has by now been extensively studied [7].

In the case of *Drosophila* it has been shown [54, 70] that flies remembered their location after their location was changed, and that this is missing in certain *Drosophila* mutants. It was pointed out [54] that the prefrontal cortex in humans and other primates similarly allows remembering the location of an object [71–73].

Likewise, we suggest that making decisions in *Drosophila*, as described in the present report (attraction to light versus repulsion by methyl eugenol), is similar to decision making in humans and other primates [1–12]. An example of such decision making in humans is the combination of a pleasant odor (jasmine) together with an unpleasant odor (indole) [74].

Putting all these things together, it appears that several processes once thought to be limited to the prefrontal cortex of “highest” animals are in fact widely distributed.

In humans the prefrontal cortex sometimes functions abnormally. This results in behavioral defects such as in patients who have had their prefrontal cortex removed [75] or in criminals who may have a deficiency in their prefrontal cortex [76]. We hope that studies like those reported in *Drosophila* will help to understand and alleviate such human problems.

## IV METHOD FOR ISOLATION OF THESE *DROSOPHILA* MUTANTS AND FOR ASSAYS USED HERE

Male *D. melanogaster* (strain Canton-S) were mutagenized with 25mM ethyl methane sulfonate using standard procedures (see Supplement) and mated to C(1)DX females. The resulting F1 males and females, between 1 to 10 days old, were tested in a tube made of three pieces cut from a 4 L graduated cylinder (Fig 14). Located at the closed (right) end of the tube were 80 lux from an LED light source and 0.1M methyl eugenol (Sigma-Aldrich, CAS No. 93-15-2) in 1.5% agar. The open (left) end was covered with parafilm and the tube was placed into a dark 34° C room. The flies, after 30 minutes in the dark in a covered dish (Fisher catalog no. 3140), were placed into the left end of the tube. Then the light was turned on. Flies which had spread away from the third of the tube nearest to the entry point were collected after 10 minutes. See Supplement for details of these procedures.

**Fig 14.**
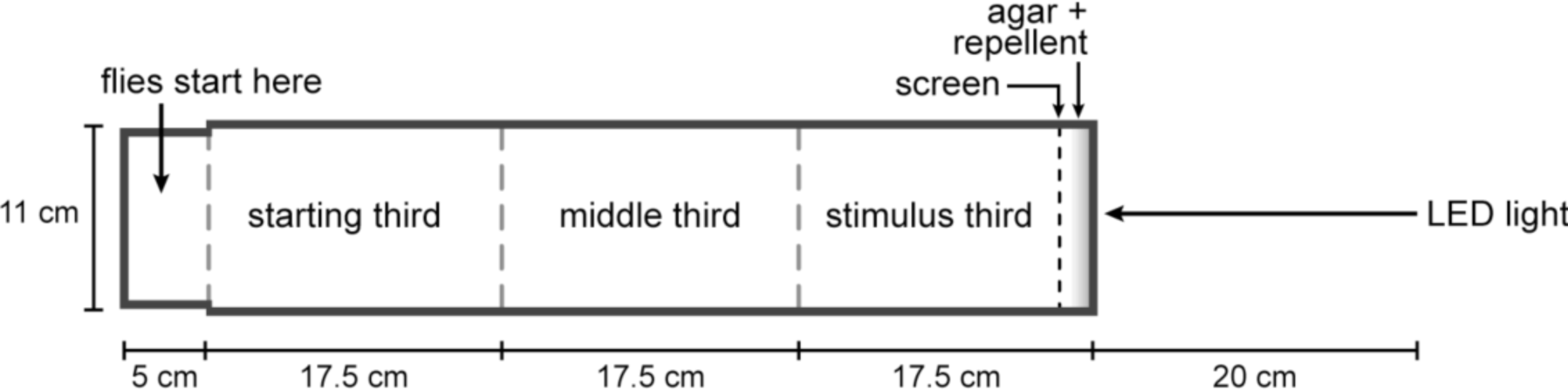
**Apparatus for isolation of mutants.** See text.

Those males that appeared mutant were then tested in a smaller setup (Fig 2), between 1 and 10 days of age. It was made from a cut test tube (18.5 x 2.5 cm, Fisher no. 14925N) connected to a shell vial (9.5 x 2.5 cm, Kimble Chase no. 60931-8) and at right to a cap (2.5 x 3 cm) cut from the bottom of shell vials. Unless otherwise noted, the cap contained 0.1M methyl eugenol in 1 ml of 1.5% agar and a fabric tulle screen was used to prevent flies coming into contact with the repellent. Flies were incubated in the shell vial for 30 minutes in the dark with a cotton plug. At 15 minutes into incubation the cap containing methyl eugenol was attached to the test tube. The open end of the tube was covered with parafilm. At 30 minutes, the parafilm was removed and the shell vial with flies was connected to the tube. Then the light was turned on for 20 minutes, during which measurements of the number of flies in each third of the tube were taken every minute. Unless otherwise noted, a fluorescent lamp was used to provide a 4000 lux light source. All characterization experiments used this small-scale test. For more details of the assay, see Supplement.

Later flies were tested in the same manner as early flies.

## ACKNOWLEDGEMENTS

We are most grateful to The Camille and Henry Dreyfus Foundation for six years of grants in support of this undergraduate research program. This project was initiated by Timmie Sharma and Julius Adler. The mutants were isolated by Timmie Sharma, Robert P. Downing, Charles G. Starr, Jay A. Doshi, Matthew N. P. Vogt, Deanna M. Luzenski, and Nathan E. Fons. The mutants were identified by Natalie M. Sciano and Nathan Eickstaedt. Further characterization of the mutants was performed by Timmie Sharma, Natalie Sciano, Robert Downing, Charles Starr, Matthew Vogt, Jay Doshi, Nathan L. Eickstaedt, Leann M. Erlien, Nathan Fons, Reuben J. Hoffmann, Hertina E. Kan, Alex R. Kleven, Lynn K. Lukoskie, Deanna Luzenski, Nathan J. Menninga, Monika Ramnarayan, Avi D. Stricker, Sarah K. Warzon, Lar L.Vang, and Andrew M. Winter. Work on other GAL4 driver lines and central complex mutants was carried out by Laura B. Burbach, Eleanor K. Ganz, and Hannah A. Hecht with the help of Sarah A. Cook, Meghan M. Cohen, Sarah C. Drewes, Nicholas A. Dykstra, Nathan Fons, Alison Heydorn, Hertina Kan, Nathan Menninga, Amelia Remiarz, Avi Stricker, and Lar Vang. Lar Vang is currently an associate research specialist in Adler’s laboratory. Robert A. Kreber, a research specialist in Barry Ganetzky’s laboratory, has helped us greatly in studies of the genetics of our mutants. Julius Adler thanks Barry Ganetzky for teaching him about fruit flies. Thanks to Millard Susman for criticisms. We are thankful to Laura Vanderploeg for the beautiful art work.

---

**S7 Fig.**
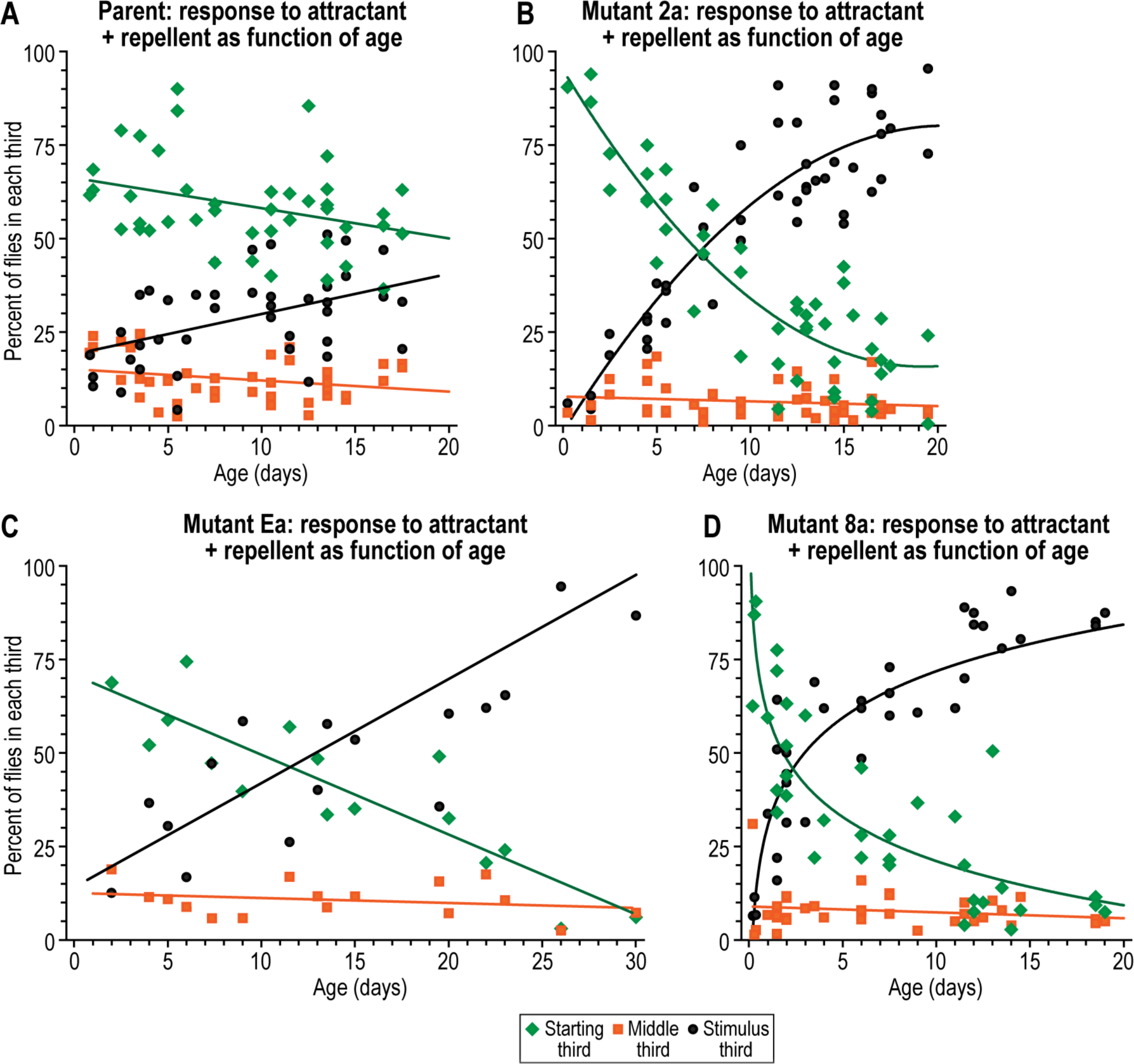
Response to attractant (light, 4000 lux) together with repellent (methyl eugenol, 0.1M) at 34°C as function of age after eclosion. (A) Parental response, (B) Mutant 2a response, (C) Mutant Ea response, and (D) Mutant 8a response. Approximately 10 to 20 flies were used per trial.

**S9 Fig.**
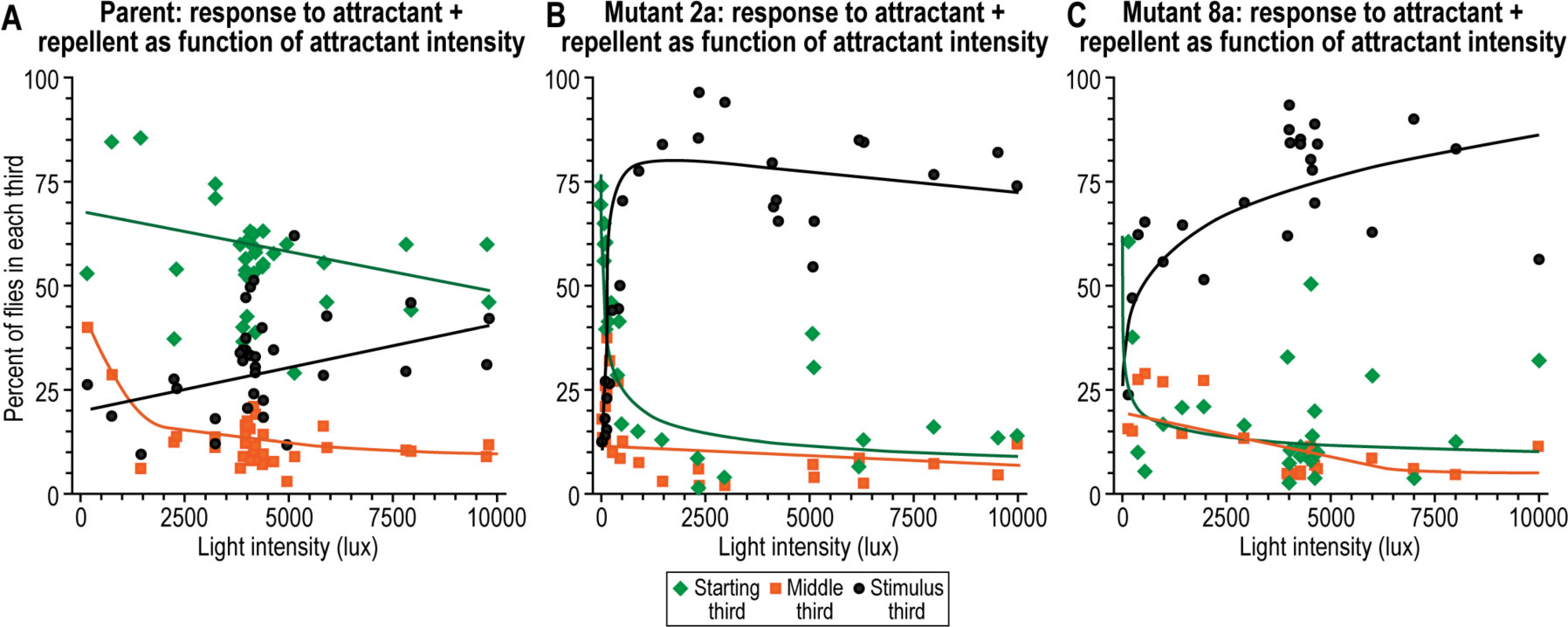
After eclosion, response to attractant (light) together with repellent (methyl eugenol, 0.1M) at 34°C as function of attractant intensity. (A) Parental response, (B) Mutant 2a response, and (C) Mutant 8a response. Flies were tested between the ages of 12 to 20 days old. Approximately 10 to 20 flies were used per trial.

**S10 Fig.**
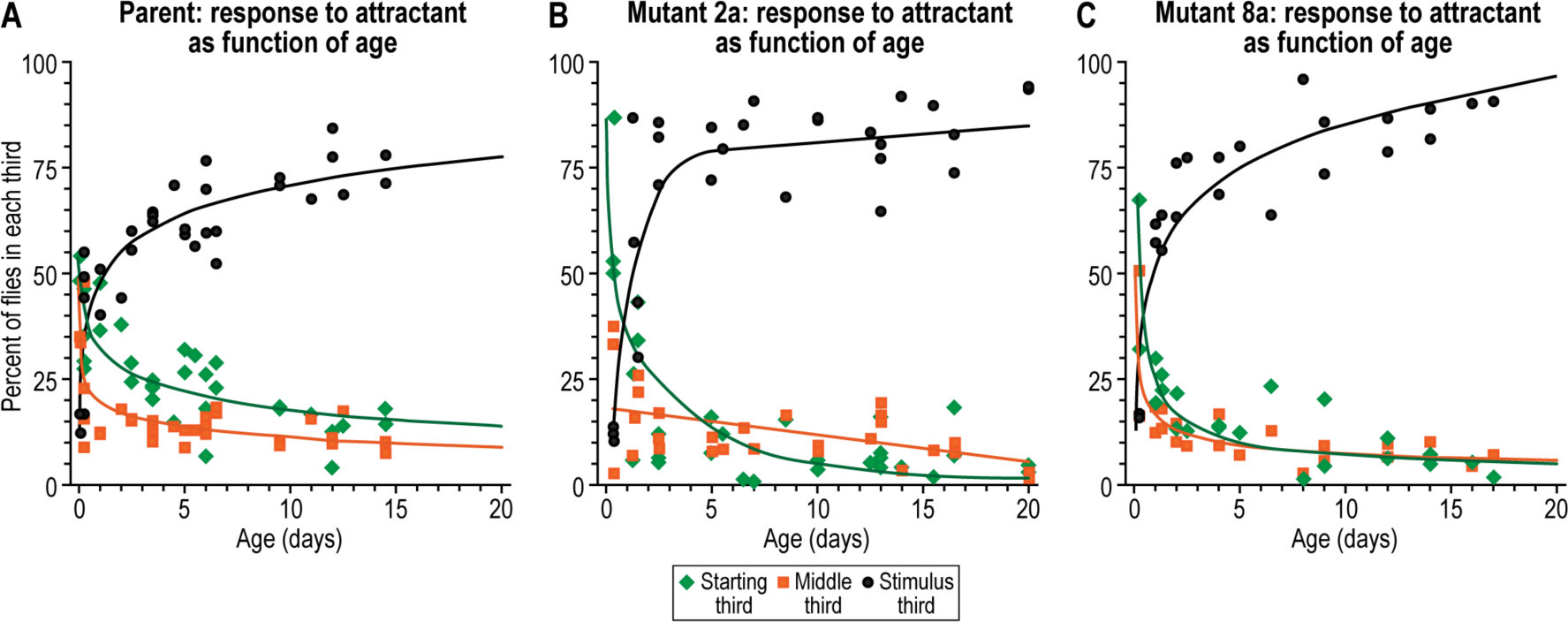
Response to attractant (light, 4000 lux) at 34°C as function of age after eclosion. (A) Parental response, (B) Mutant 2a response, and (C) Mutant 8a response. Approximately 10 to 20 flies were used per trial.

**S11 Fig.**
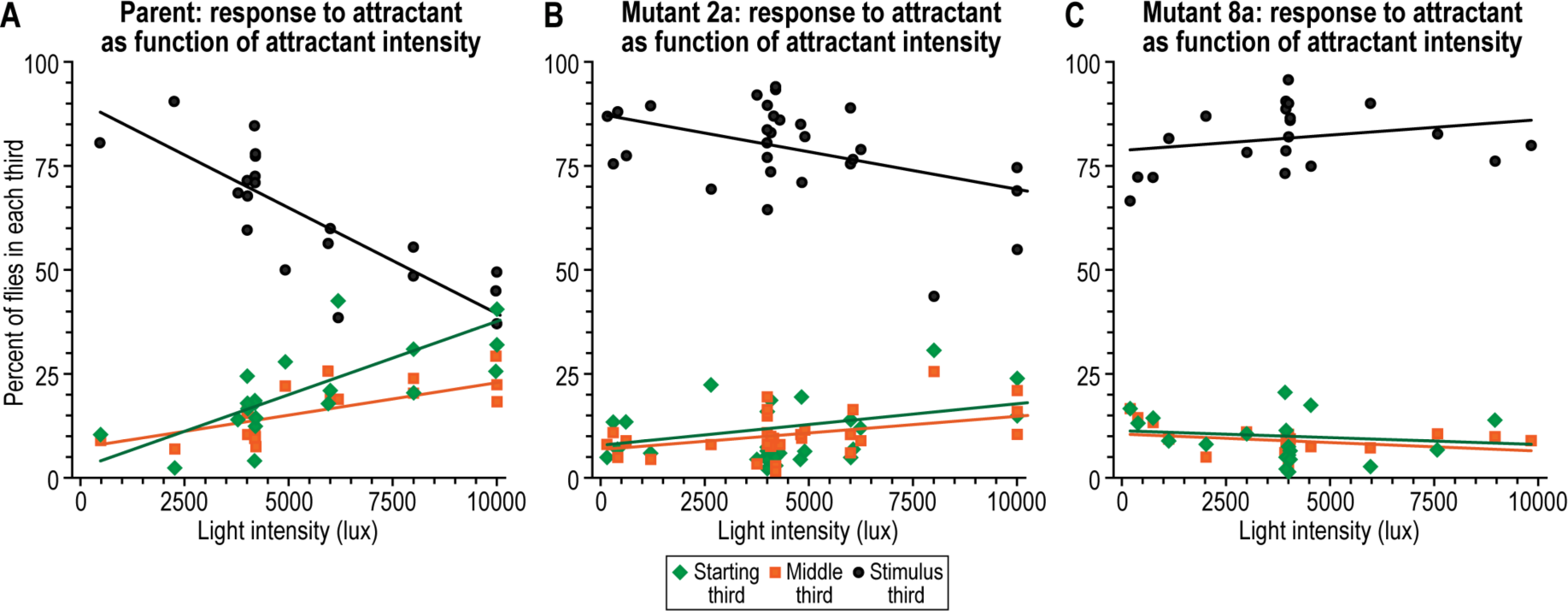
After eclosion, response to attractant (light) at 34°C as function of attractant intensity. (A) Parental response, (B) Mutant 2a response, and (C) Mutant 8a response. Flies were tested between the ages of 10 to 20 days old. Approximately 10 to
20 flies were used per trial.

**S12 Fig.**
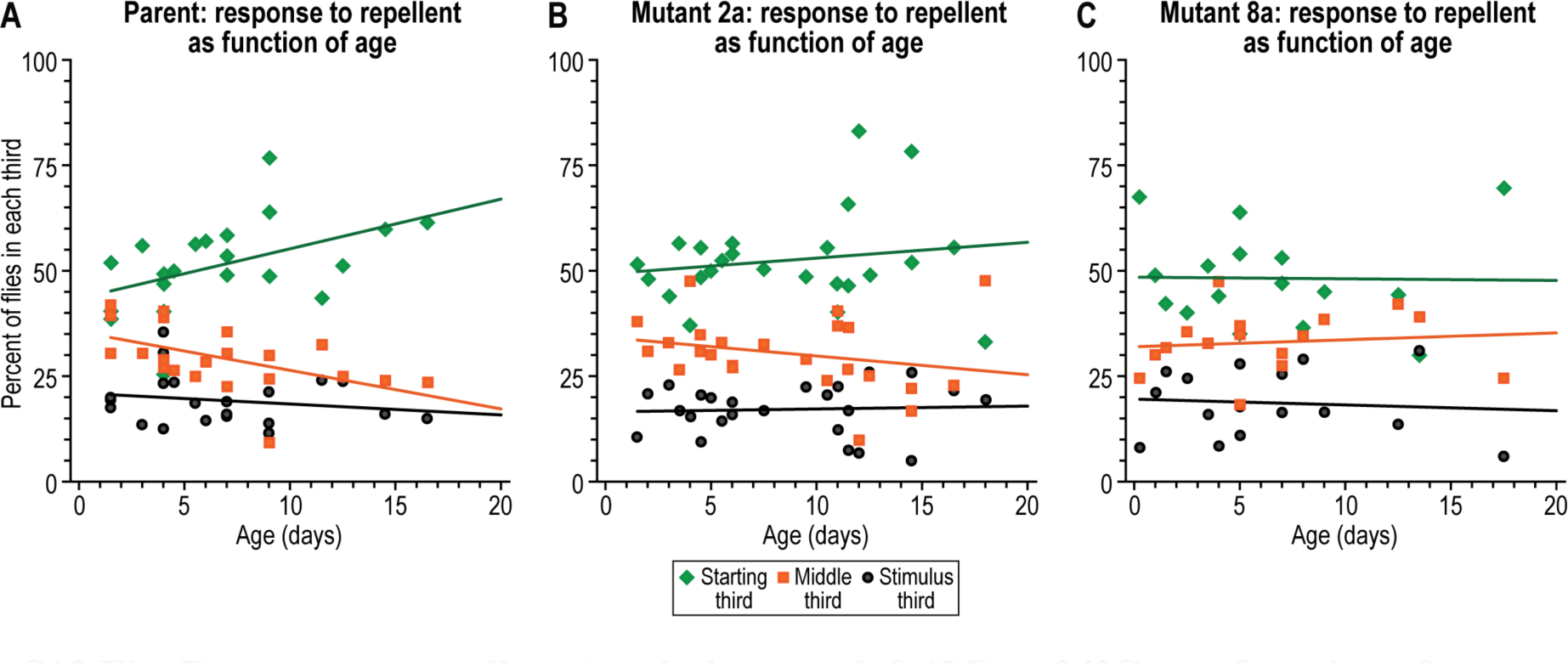
Response to repellent (methyl eugenol, 0.1M) at 34°C as a function of age after eclosion. (A) Parental response, (B) Mutant 2a response, and (C) Mutant 8a response. Approximately 10 to 20 flies were used per trial.

**S13 Fig.**
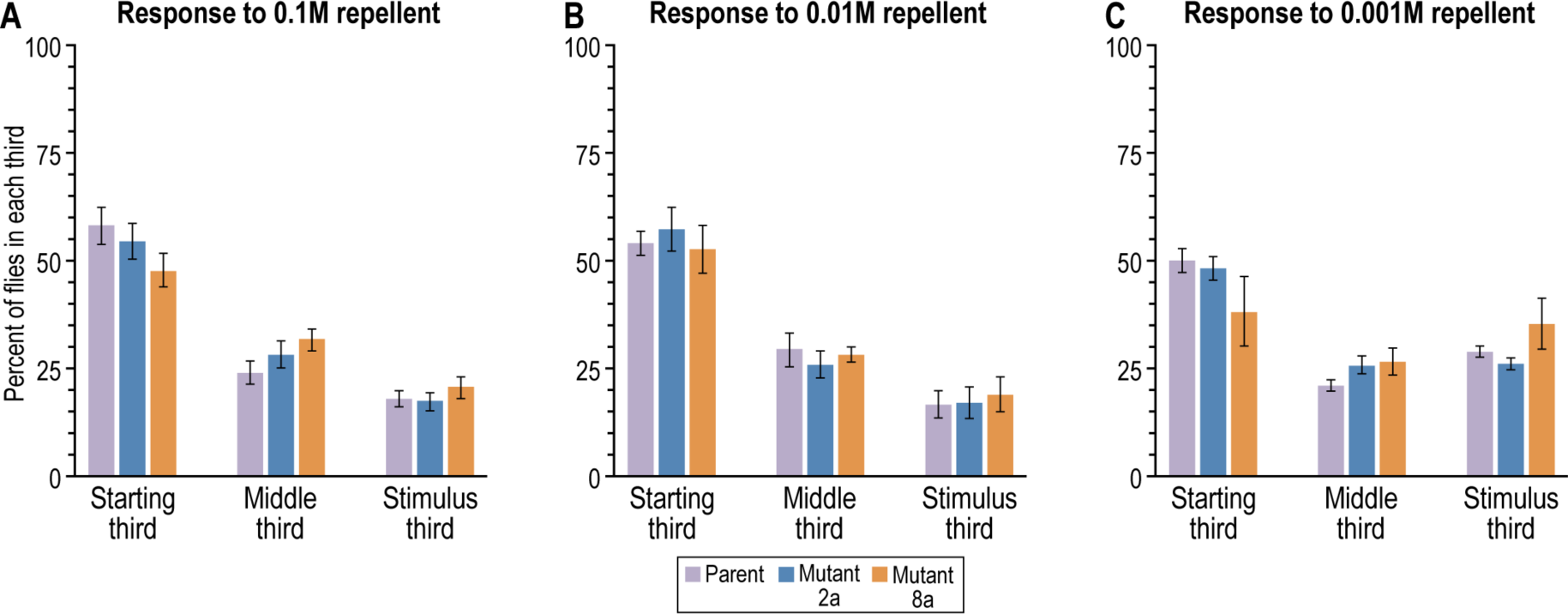
Response to repellent (methyl eugenol) as a function of repellent concentration. (A) Response to 0.1M repellent; parent (n=7), Mutant 2a (n=12), and Mutant 8a (n=7). (B) Response to 0.01M repellent; parent (n=4), Mutant 2a (n=3), and Mutant 8a (n=4). (C) Response to 0.001M repellent; parent (n=3), Mutant 2a (n=3), and Mutant 8a (n=3). Flies were tested between the ages of 9 to 20 days old. Approximately 10 to 20 flies were used per trial. Data are mean±SEM.

**S15 Fig.**
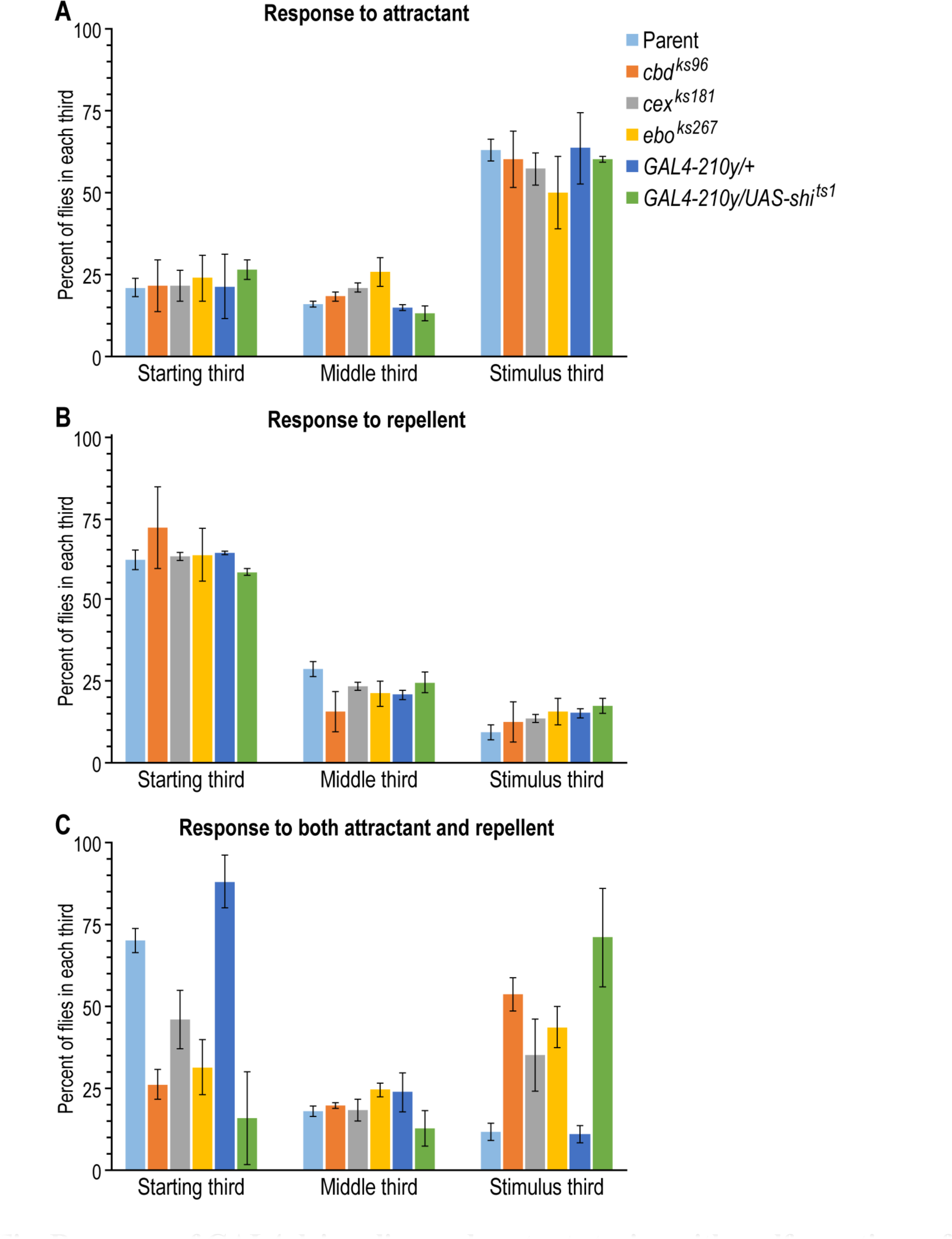
Response of GAL4 driver line and mutant strains with malformations of the central complex. (A) Response to attractant (light, 1000 lux) only; parent (n=7), *cbd*^*ks96*^ (n=3), *cex*^*ks181*^ (n=3), *ebo*^*ks267*^ (n=3), *GAL4-210y/*+ (n=3), *GAL4-210y/UAS-shi*^*ts1*^ (n=3). (B) Response to repellent only (0.1M); parent (n=7), *cbd*^*ks96*^ (n=3), *cex*^*ks181*^ (n=3), *ebo*^*ks267*^ (n=4), *GAL4-210y/*+ (n=4), *GAL4-210y/UAS-shi*^*ts1*^ (n=3). (C) Response to attractant (light, 1000 lux) together with repellent (0.1M methyl eugenol); parent (n=8), *cbd*^*ks96*^ (n=6), *cex*^*ks181*^ (n=5), *ebo*^*ks267*^ (n=4), *GAL4-210y/*+ (n=4), *GAL4-210y/UAS-shi*^*ts1*^ (n=5). Flies were tested at 34°C and between the ages of 3 to 14 days old. Approximately 10 to 20 flies were used per trial. Data are mean±SEM.

## SUPPLEMENTAL METHODS

The attractant used here, light, has been extensively studied in *D. melanogaster* [1, 2]. Early steps of vision are now well known, both in vertebrates and invertebrates [1, 3], while later steps for vision are being investigated [4]. Such is true also for response to smell, both in vertebrates and invertebrates, for the earliest steps [5–7] and also for later steps [8].

The repellent used here was methyl eugenol (also named 4-allyl-1,2-dimethoxy-benzene, obtained from Aldrich, Milwaukee) [9–12]. Various repellents other than methyl eugenol were tried by us (benzaldehyde, ethyl hexanoate, trans-2-hexenal, methyl salicilate, 2-phenyl ethanol, salicilate) but only methyl eugenol was found potent enough to bring about the repulsion needed when this amount of light was used. (Methyl eugenol appears to be an attractant for *D. melanogaster* at lower concentrations like 10^−5^ to 10^−6^ M, unpublished data, of Adler and Vang) Methyl eugenol has not previously been used as a stimulus in *D. melanogaster* to the best of our knowledge, but a hundred years ago the Indian biologist Frank Howlett discovered that methyl eugenol is a powerful attractant for a different, tropical fruit fly [9–12] and likely that it also is a repellent at higher concentrations [9].

*D. melanogaster* (strain Canton-S) were maintained on standard cornmeal-molasses agar medium at room temperature (21-23°C) in a room that was light for 12 hours and dark for 12 hours. Male flies were mutagenized with 25mM of the mutagen ethyl methane sulfonate added to 1% sucrose by Robert Kreber in Barry Ganetzky's laboratory [13]. These were mated with virgin C(1)DX attached-X females. Then after fertilization and egg laying the males and females were removed at about six days, then at about two weeks adults appeared (the F1 generation). About 1,500 of these adults were tested as follows between the ages of 1 and 10 days.

A 4-liter graduated cylinder (Nalgene polymethylpentene, Fisher catalog no. 3663), 58 cm long and 11 cm wide, was cut at 5.5 cm to reject the spout and then cut to provide three parts each 17.5 cm in length for isolation of mutants. Holes smaller than the size of *Drosophila* were placed all over to allow exchange of gasses (O_2_, CO_2_, etc.) (In later procedures holes for gas exchange were not used.) Fig 14 shows the three parts, called "stimulus third" for locating the attractant and repellent, “middle third”, and “starting third” for being next to a dish where the flies start out.

About 500 F1 flies (males together with females) were put into a dish (Pyrex crystallizing dish, Fisher catalog no. 3140) through a hole in a piece of cardboard that covered the dish, a book was then placed over it for weight, then at −30 minutes all this was placed into a dark 34°C room. Twenty mL of 1.5% agar (that had been melted and then kept at 65°C) was mixed by vortexing with 34 uL (1x10-2 M final concentration) of methyl eugenol for 10 seconds. This was pipetted vertically into the stimulus third at room temperature and then covered with a screen 5 cm away to prevent flies from getting killed by the repellent. The middle third at room temperature was then attached to the stimulus third by means of 48 cm-wide transparent packaging tape (Scotch, from 3M Stationery Products Division, St. Paul) and the other end of the middle third was covered with Parafilm. Then the combined stimulus third and middle third were placed into the dark 34°C room at −20 minutes to allow heating and to allow diffusion of the repellent. At −5 minutes the Parafilm was removed and the starting third (previously warmed at 34°C with Parafilm at its left end) at its right end was attached by means of packaging tape to allow a further diffusion of the repellent. At zero time the Parafilm was removed and the dish containing the flies (cardboard and book at this time removed) was horizontally added by pushing it into the starting third. (The dish had been provided with enough 1.9 cm white-label tape, Fisher catalog no. 15-938, to make it fit into the slightly larger starting third.) Then the light source was turned on. It was a white LED light set at 8.6V placed 20 cm away from the stimulus third where the light intensity was measured as 80 lux. (Incidentally, these attached-X females are not attracted to the light at 80 lux in presence of methyl eugenol at 1x10^−2^ to 1×10^−1^M but they are attracted at 4000 lux in presence of these amounts of methyl eugenol; the male flies, on the other hand, are not attracted to light under these conditions, nor are standard females). To allow better visibility throughout the procedure, a 45 cm fluorescent light (15 watts, cool white, lamp model VTU15RTCPCO used without plastic cover, American Fluorescent Corp., Waukegan, Il, or local hardware stores such as Mennards, Madison, WI) was placed perpendicularly across the room behind a black curtain. (Instead of using the LED light, an alternative procedure was to use a 45cm fluorescent light at 4000 lux.) At 10 minutes the test was ended, and then flies in the middle third (about 5% of the total) together with those in the stimulus third (about 3% of the total) were combined, and the males there were studied further as described next.

The males from above were then assayed on a small scale as in Fig 2, between 1 and 10 days old. It is like Fig 14 except that there was no dish and it was smaller – 31 cm in length and 2.5 cm wide. At the stimulus end was a 3 cm piece (cut from the closed end of a shell vial, Fisher catalog no. 03-339-30J) containing at its end 1 ml agar plus repellent (0.1M methyl eugenol) and covered with a 6 cm × 6 cm white-cloth screen (“tulle fabric” from Gifts International Inc. www.giftsintl.com or local fabric stores such as Hancock Fabrics, Madison) (or a metal screen); in the middle was an 18.5 cm piece cut from a 20 cm × 2.5 cm test tube (Fisher catalog no. 14925N); at the start was a 9.5 cm piece (a shell vial) for housing the flies at the beginning. The three pieces were put together by means of two 2 cm adapters (cut from a Beckman plastic centrifuge tube, Spinco catalog no. 344058). Illumination was with the fluorescent light source placed perpendicularly and 5 cm away from the agar end, where the light intensity was measured at about 1000 or 4000 lux.

